# *Heteractis magnifica* Sea Anemone Population Dynamics at Village Reef, Perhentian Kecil, Malaysia: Monitoring of Growth, Formations, and Hosting Status

**DOI:** 10.1101/2022.02.04.479141

**Authors:** Melissa Versteeg, Alanah Campbell, Hidayah Halid

## Abstract

The coastal waters of Malaysia have been known to allow proliferation of sea anemone assemblages, which are resident species of tropical coral reefs. Along the Perhentian Islands of Terengganu, no efforts have been made thus far to investigate the presence and population dynamics of sea anemone assemblages locally. In this study, *Heteractis magnifica* assemblages at Village Reef at Perhentian Kecil were monitored during May, July, and August of 2020, thus providing a first assessment of their abundance. Sea anemone formation size, individual size, habitat location, and hosting status of anemonefish were assessed. Results demonstrate significantly larger counts of individuals within aggregated formations in the patch reef as compared to the fringe reef, without the presence of larger individual sizes. In addition, *Heteractis magnifica* specimens that were actively hosting anemonefish had significantly larger cover, larger individual sizes, and demonstrated higher individual counts within their aggregated formations compared to non-hosting specimens. There was no significant overall effect of time on sea anemone growth throughout the monitoring period, nor were there any significant changes in abundance levels regarding formation make-up throughout the monitoring phase. However, time related effects where present upon data inspection per assessment period. The restricted time frame of the monitoring period could play a role in explaining the absence of overall time related effects. A prolonged monitoring will help to further understand *Heteractis magnifica* population dynamics at this location, which can increase knowledge capacity for reef management and conservation strategies.

## INTRODUCTION

The sea anemones’ (*Actiniaria)* ability to reproduce asexually, and their lack of skeletal structure allows rapid formation and community expansion in suitable environments (Steinberg *et al*. 2020). Sea anemones can continually produce viable nodules for colonisation of neighbouring patches, where, given favourable environmental factors, sexual reproduction is actively suppressed in favour of rapid, asexual colonisation (Brace & Quicke 1986). Environmental parameters that influence sea anemone abundance and growth include temperature, seasonal effects, radiance and nutrient loadings, pollutants, and depth (Brolund *et al*. 2004; Chomsky *et al*. 2004; Holbrook & Schmitt 2005; Muller-Parker & Davy 2001; Thomas *et al*. 2014). Furthermore, when in the presence of live coral, sea anemones have been found to demonstrate higher expansion rates and will utilize aggressive strategies towards neighbouring corals during competition over suitable substrate (Liu *et al*. 2009).

To the east of peninsular Malaysia, the coral reef habitats of the Perhentian Islands have undergone dramatic changes in previous years (Islam *et al*. 2013). Longitudinal monitoring demonstrates a profound drop in live coral cover, with the 2008 averages calculated at 50.74%, whilst a mere 35.50% of the reefs contained live coral cover by 2019 (Reef Check Malaysia 2008; Reef Check Malaysia 2018). Furthermore, a sharp rise in tourism (Department of Marine Park Malaysia 2016) with subsequent increases in infrastructure development, tourism impacts, and changes to water quality, are negatively influencing the environmental quality of the Perhentian Islands (Nasir *et al*. 2017; Kurniawam *et al*. 2016). Although research focusses on monitoring changes in coral states throughout the Perhentian Islands, no studies to date are monitoring changes in sea anemone distribution patterns.

To protect Perhentian coral reefs from degradation and to stimulate coral recovery, the Perhentian Islands were gazetted as a Marine Park (MP) in 1994 (Department of Marine Park Malaysia 2014). At the MP, regulations are designed to protect essential marine ecosystems so as to ensure sustainability of vital fish stocks and marine environments (Department of Marine Parks Malaysia 2016). Government departments, local businesses, and social enterprises alike engage in reef restorative activities such as coral planting projects, promotion of ‘reef friendly’ tourism techniques, removal of toxic or smothering materials, and educational outreach. Despite these efforts, large-scale effectiveness is significantly impacted by restraints in staff capacity, lack of adequate financial resources, and logistical limitations (Islam *et al*. 2013). At the Perhentian Islands, littering, overfishing, and discarded fishing gear are degrading the reef states and monitoring studies continue to present trends of declining coral health (Reef Check Malaysia 2008-2018). As sea anemones require environmental parameters similar to that of corals, the changing environment may influence the hosting sea anemones’ opportunity for colonisation and growth (Tkachencko *et al* 2007). In environments additionally marked by high sedimentation, and water temperatures outside the thermal ranges suited to algae proliferation, sea anemones increase growth rates, asexual reproductive rates, and increase aggressive attacks on neighbouring corals (Chomsky *et al*. 2004; Liu *et al*. 2009; Liu *et al*. 2015).

The clear shallow waters that serve as primary habitat to tropical corals and sea anemones are characteristically low in available nutrient levels (Hopley 2011; Mohamed *et al*. 2019; Schwartz 2005), with dependent species relying on high nutrient recycling adaptations, storage mechanisms, and a capacity for efficient nutrient capture in the water column (Ortega *et al*. 1988; Savage 2019). At low nutrient levels, corals, sea anemones, and algae coexist. Upon altering dissolved P and N levels however, imbalance between these three groups sets in (Liu *et al*. 2009). Initially, high nutrient levels are catalytic to higher chlorophyll densities, with increases in energy availability and growth rates for corals (Savage 2019). Further rises however are a determinant for increased algae and sea anemone abundance.

Nutrient dispersal along Malaysia’s coastal waters is biologically managed through a suit of mechanisms including mixing, upwelling, currents, tides and drifts, terrestrial run-off and seasonal climatic patterns including monsoon events. Research on global nutrient flow mechanisms demonstrate that water movement allows deeper lying nutrients to be transported to shallow regions, via currents or drainage from terrestrial water systems (McPhee-Shaw *et al*. 2007; Mohamed *et al*. 2019; Powley *et al*. 2017; Sardessai *et al*. 2007). The supply of nutrients to the Perhentian Islands corresponds with water displacement events occurring during the seasonal cycles of the Northeast monsoon, and as such, marine species like hosting sea anemones, which rely on these nutrients for growth, may expand coverage in synchrony with the Northeast monsoon. Rivers draining into the South China Sea along Terengganu predominantly regulate nutrient supply to the Perhentian Islands, with highest concentrations found during the post monsoon phase (Adiana *et al*. 2014; Mohamed & Amil 2015). Along the Perhentian reefs, distributions of dissolved NO3 and PO4 are between 16 to 83 times higher during the post monsoon phase, with the greatest depletion levels located at depths of three meters, and a maximum concentration between three- and six-meters depth (Mohamed *et al*. 2019). Besides delivering required nutrients, discharge of pollutants into river systems can also occur and results in elevated values of trace metals in neighbouring coastal regions (Shazili *et al*. 2006).

Experiments also reveal high survival rates when sea anemone specimens are split into sections, demonstrating their capacity to propagate through axial thinning and tearing of tissue via longitudinal fission (Porat & Chadwick-Furman 2004; Scott *et al*. 2014). Their ability to reproduce via such mechanisms displays adaptivity to challenging environments and highlights mechanisms of competitive re-colonisation. In the case of clustered sea anemones, asexual reproductive techniques, including longitudinal fissure, are proposed to underly aggregate formations as physically touching sea anemones are reported to constitute clones (Allen 1975; Dunn 1977).

Sea anemones with hosting capacity additionally undertake an obligatory symbiosis with anemonefish (*Amphiprion spp*.) for nutrient intake, including a direct transfer from symbiont to host (Cleveland *et al*. 2011; Norin *et al*. 2018; Porat & ChadwickFurman 2004; Roopin & Chadwick 2009). As such, actively hosting sea anemones show increased growth rates and regeneration rates compared to their non-hosting counterparts (Holbrook & Schmitt 2005). Moreover, actively hosting sea anemones experience increased oxygenation, resulting in higher respiratory and growth rates compared to sea anemones without symbiotic anemonefish residents (Herbert *et al*. 2017; Szcezebak *et al*. 2013) and the distribution patterns of hosting sea anemones are tightly linked to successful recruitment of symbiotic anemonefish (Elliot & Mariscal 2001). Furthermore, their hosting capacity allows sea anemones increased odds for successful recovery following bleaching events (Norin *et al*. 2018). With increases in sea temperature and subsequent rises in bleaching risk, hosting sea anemones maintain a higher likelihood of adaptation and survival under adverse conditions (Pryor *et al*. 2020).

Within the Malaysian waters, large formations of sea anemone aggregates or clustered formations can be found naturally (Dunn 1977), including sea anemone species with hosting capacity. In Terengganu, on Perhentian Kecil, a reef site adjacent to the town village, called ‘Village Reef’ displays large aggregates of hosting sea anemones. Proximity of the reef site to the village indicates a possibility of anthropogenic effects, which have been found to decrease hard coral cover in addition to affecting coral community composition (Crehan *et al*. 2019). If the reefs of Perhentian Kecil experience environmental settings conducive to hosting sea anemone proliferation, we expect the local sea anemone aggregates to expand whilst favourable conditions remain in play.

As previously mentioned, at Village Reef, large assemblages of sea anemone species *Heteractis magnifica* are present (**Figure 1**). To date, no research has been conducted to investigate *Heteractis magnifica* population dynamics in this specific region via quantitative study. As such, the current study sought to monitor the assemblages of *Heteractis magnifica* at Village Reef so as to develop a baseline measurement of their abundance levels, formation patterns, and hosting status, in addition to examining *Heteractis magnifica* growth.

**Figure 1.**
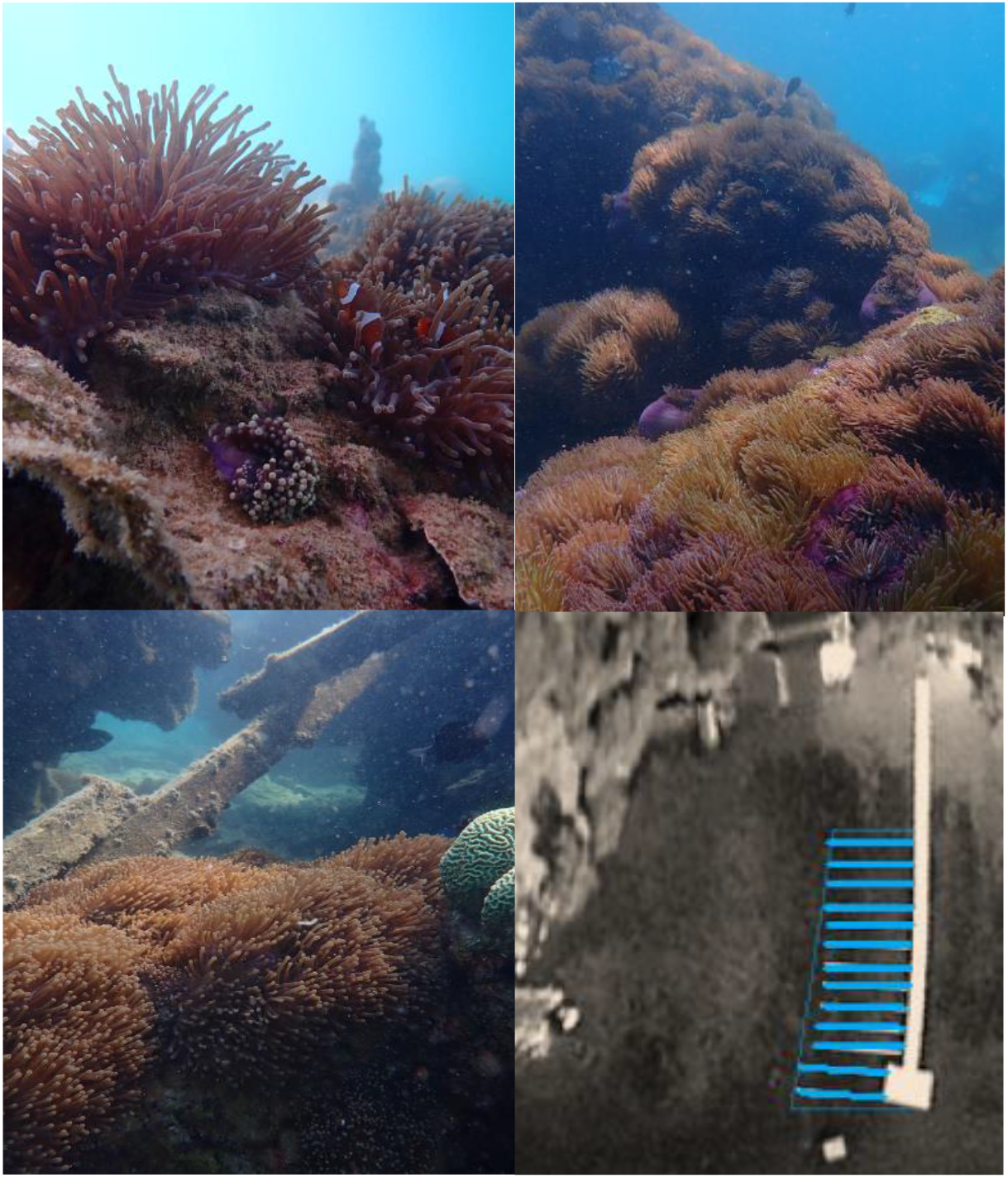
Images of Heteractis magnifica formations at research site Village Reef, including a schematic of study site area (bottom right).

The current study expects to find time related effects on sea anemone size and cluster formations where formation cover, individual size, and clustered counts increase over time, by reasoning that synchrony with post Northeast monsoon nutrient availability will stimulate growth at the reef site. In addition, a significant difference between reef habitats is expected, with the patch reef deemed more favourable for *Heteractis magnifica* proliferation based on depth parameters. In the patch region at Village Reef, we expect higher *Heteractis magnifica* abundance levels and increased formation size, individual size, and cluster counts over time, compared to the fringe reef. More so, as active hosting status is correlated to increased growth and reproduction rates (Holbrook & Schmitt 2005; Porat & Chadwick-Furman 2004), we also expect the actively hosting sea anemones at Village Reef to be significantly larger than their non-hosting counterparts, including larger individual sizes over time. Finally, with previous studies highlighting an important role for asexual reproduction under favourable conditions, (Scott *et al*. 2014; Brace & Quicke 1986) the current study expects increased clustered formations over time.

This study offers a first examination of localised *Heteractis magnifica* formations. Insights serve to increase local distribution knowledge and informs reef management and conservation. In addition, resultant knowledge may aid coral planting strategies at the Perhentian Islands by expanding the ability with which to identify temporal environmental patterns that favour expansion of hosting sea anemones over that of local corals.

## MATERIALS AND METHODS

### Study Area and Study Population

Data was collected at the Village Reef survey site on the South East of Perhentian Kecil (central coordinates: 5°53’39.05”N, 102°43’37.61”E). Village Reef comprises of a fringing, coral dominant reef section located in the lower intertidal zone, and a deeper region of patch reef. Throughout the shallower fringing reef, the effects of the semi-diurnal tides can create partial exposure during low tides, making parts of this habitat unsuited to *Heteractis magnifica* (Fautin 1991; Muller-Parker & Davy 2001). The patch reef displays dead coral bommie structures which have since been recolonized by stony and soft corals, sponges, algae species, and sea anemones.

Within the site boundaries, two hosting sea anemone species were identified: *Stichodactyla gigantea* and *Heteractis magnifica*. Only two counts of *Stichodactyla gigantea* were present during August 2020, compared to 224 formations of the *Heteractis magnifica* at that time. Interspecies differences in reproductive strategies (Aubert 2014), habitat parameters (Elliot & Mariscal 2001), and size (Fautin & Allen 1992), determined for the exclusion of the *Stichodactyla gigantea* specimens to control for confounding effects.

### Data Collection Method

Between May and August of 2020, abundance, size, hosting status, and formation markers of hosting sea anemone species *Heteractis magnifica* were monitored using SCUBA. All data collection sessions took place between 8.30am and 11.59am, and visibility had to be over five meters as a prerequisite to diving. Within the survey area, ten 20 meter transects were laid out in parallel using a 225° south-west bearing, in addition to cross-referencing from a stable landmark (see **Figure 1**). Distance between parallel transects was set at 4 meters to allow accurate monitoring without overlap. Following transect placement, two research divers regressed along the transect line, keeping a two-meter width perpendicular to the transect. Each *Heteractis magnifica* formation within the survey area was identified, measured, and observed to record the variables under study: reef habitat, hosting status, formation size and cluster counts.

### Reef Habitat

Reef habitat was categorised as ‘patch’ or ‘fringe’ per 80m^2^ transect, based on in-situ depth and reefscape observations. Fringing reef areas were marked by shallower depth <1.5m, absence of coral bommies, and significantly higher levels of live coral cover, where interconnected carpets of hard corals dominated the sea floor. Of the total survey site, 480m^2^ was defined as fringing reef, which comprised of five transects measuring the shallowest regions of the research site. At Village Reef’s fringing reef, 20 *Heteractis magnifica* formations were recorded during August 2020, with a cumulative cover of ∼2.4m^2^.

The shallowest three fringing reef transects contained three solitary *Heteractis magnifica* formations within 240m^2^, representing 0.45% of the total abundance during August. As this area skewed the data distribution significantly, and literature indicated unfavourable habitat requirements, these transects were excluded as outliers from the final analysis. The remaining transects covered a survey area of 560m^2^, of which 240m^2^ contained fringe reef.

Patch reef was marked by depth >1.5m, with a maximum, tide-dependent depth between 5-6 meters. The patch reef housed coral bommie skeletal structures, interspersed with sandy areas, rocky substrates, and coral rubble. Four transects were located within the patch reef of the survey site at Village Reef, with a combined area of 320m^2^. In August, 224 *Heteractis magnifica* formations were recorded on the patch reef, with a cumulative cover of ∼34.5m^2^.

### Hosting Status

Actively hosting or non-hosting status was recorded, as previous results indicate effects of size and growth (Herbert *et al*. 2017; Elliot & Mariscal 2001). *Heteractis magnifica* specimens were coded as actively hosting if they were inhabited by symbiotic anemonefish. The temporal nature of symbiosis between hosting sea anemones and other fish species such as the *Dascyllus trimaculatus* (Fautin & Allen 1992), made that only anemonefish species were included in hosting assessments.

Upon encountering a *Heteractis magnifica* specimen, indicators of hosting status were observed, including clear presence of anemonefish, or indirect indicators such as irregular movements between tentacles plus partial observations of anemonefish. If during in-situ observations a given sea anemone appeared not to be actively hosting, they were inspected more closely to check for juvenile species. If active hosting was suspected but not confirmed, video footage was recorded and reviewed to inform hosting status ex-situ.

### Heteractis Magnifica Formations

Formations were examined three times between May and August 2020. Sea anemones dwelling in clustered formations are found to have higher growth rates, higher reproductive success, and can harbour more symbiotic anemonefish species, which in turn promotes increased nutrient deposition, and higher levels of host oxygenation (Cleveland *et al*. 2011; Herbert *et al*. 2017; Holbrook & Schmitt 2005; Norin *et al*. 2018; Porat &Chadwick-Furman 2004). Specimens were marked as clustered formations when a fully expanded individual’s tentacles could touch a neighbouring sea anemone (Allen 1975). In the event of ambiguity, video recordings were made for ex-situ examination by both researchers.

### Sea Anemone Size and Clustered Individual Counts

To estimate size, a long and short axis measurement of the oral disc was taken, using a tailor’s tape, and subsequently used to conduct calculations (Hirose 1985). If the *Heteractis magnifica* was retracted, time was given for the animal to resume expansion before resuming measuring. For sea anemones present as clustered formations, the same method was applied, but using the centre of the cluster as a mid-point for axial measurements. In the event that clusters did not fully cover the substrate, or if clusters did not assume a circular or elliptical shape, an area cover estimate was recorded to adjust calculation (0-100%, estimated in increments of 10).

To calculate individual size estimates for clustered sea anemones, cluster categories were marked (Allen 1975). Categories include solitary, less than 5, less than 10, less than 15, with subsequent increments of 5 until less than 35, which was the largest cluster formation category encountered at Village Reef. The midpoint of each category was subsequently used to estimate a median number of individuals per cluster, with the minimum and maximum count per category used to determine ranges of individuals within a cluster.

To calculate individual size, the total cluster size was divided by the median number of individuals, based on methods used by other researchers (Holbrook & Schmitt 2005). For example, in a cluster of <15 individuals with a collective size estimate of 0.150m2, the mean individual size was calculated by taking the mid-point of the cluster category, in this case 12 (cluster category <15 indicates 10-14 individuals) making the individual size estimate 0.0130m2 for all individuals.

### Statistical Analysis

Data descriptive statistics and statistical analyses were run using Statistical Package for Social Sciences (SPSS) version 27.0. Time logs, depth readings, date stamps, and database inputs were all completed immediately following dives, and an interobserver analysis revealed a recording accuracy of 96,7% on all monitored variables.

For all significance tests, a threshold of .05 or was used to determine rejection or acceptance of the null hypothesis. Exploration of the dataset in SPSS showed nonnormal data distribution, thus requiring non-parametric testing of our hypotheses. In addition, as fringe reef and patch reef demonstrated substantial habitat differences, the analyses pertaining to hosting status and clustered formations were run exclusively for the patch reef.

## RESULTS

In total, 560m2 of the reefscape at Village Reef was monitored and analysed, and 640 *Heteractis magnifica* formations were recorded. In the fringe reef area 65 formations were analysed, with 575 formations analysed within the patch reef. Of the sea anemones, 77.03% were actively hosting symbiotic *Amphiprion spp*. (N=493). Of these, 65.8% had resident *Amphiprion ocellaris* (N=421), 10.9% were hosting *Amphiprion perideraion* (N=70), and two formations were hosting both anemonefish species. On average, over the entire monitoring period, 48.91% of the analysed specimens regarded solitary formations of *Heteractis magnifica*, with the remainder concerning clustered formations.

Throughout the research site, an average of 3.79 individual sea anemones were clustered together (SD=4.75), with a maximum of 32 individuals clustered within a formation. During May, the collective cover of the *Heteractis magnifica* population was 31.84m^2^, reaching 37.63m^2^ by July, and reducing to 36.88m^2^ by August. Further descriptive statistics for the study area at Village Reef are presented in **Table 1**.

**Table 1.** Descriptive statistics for Heteractis magnifica monitored within the entire research site at Village Reef, including formation and individual size, individual counts, and hosting status.

| Monitoring period | N | Formation size (m<sup>2</sup>) |  | Individuals per cluster (count) |  | Individual size (m<sup>2</sup>) |  | Actively hosting | Solitary formations | Minimum number of individuals (count) | Maximum number of individuals (count) |
| --- | --- | --- | --- | --- | --- | --- | --- | --- | --- | --- | --- |
|  |  | Mean | SD | Mean | SD | Mean | SD | (%) | (%) |  |  |
| May | 197 | .162 | .211 | 3.51 | 4.56 | .049 | .031 | 74.11 | 47.72 | 548 | 836 |
| July | 219 | .172 | .242 | 4.11 | 5.00 | .044 | .026 | 79.45 | 47.95 | 729 | 1073 |
| Aug | 224 | .164 | .249 | 3.73 | 4.90 | .047 | .036 | 77.23 | 50.89 | 676 | 992 |
| Total | 640 | .166 | .235 | 3.79 | 4.75 | .046 | .031 | 77.03 | 48.91 | 1953 | 2901 |

In the fringe reef, 73.8% of the formations contained actively hosting *Heteractis magnifica* formations (N=65). In addition, 54.48% of the sea anemones presented as solitary formations, with clustered formations containing 2.51 individuals on average. The maximum clustered formation within this reef region consisted of 12 individuals. During May, the collective cover of the sea anemones located in the fringing region at Village Reef was 1.80m^2^, which increased to 3.64m^2^ by July, and came to 2.39m^2^ by August. **Table 2** displays further descriptive statistics for Village Reef’s fringe reef.

**Table 2.** Descriptive statistics for Heteractis magnifica monitored within the fringing reef region of the research site at Village Reef, Perhentian Kecil, including formation and individual size, individual counts, and hosting status.

| Monitoring period | N | Formation size (m<sup>2</sup>) |  | Individuals per cluster (count) |  | Individual size (m<sup>2</sup>) |  | Actively hosting | Solitary formations |
| --- | --- | --- | --- | --- | --- | --- | --- | --- | --- |
|  |  | Mean | SD | Mean | SD | Mean | SD | (%) | (%) |
| May | 20 | .090 | .078 | .056 | .042 | 1.80 | 1.51 | 75.00 | 70.00 |
| July | 24 | .151 | .152 | .060 | .042 | 2.96 | 2.74 | 75.00 | 45.83 |
| Aug | 21 | .114 | .113 | .047 | .034 | 2.67 | 2.60 | 71.43 | 47.62 |
| Total | 65 | .118 | .114 | .054 | .039 | 2.48 | 2.28 | 73.81 | 54.48 |

At Village Reef’s patch reef, just under half of the 575 formations contained solitary formations, at 48.35%. Of the clustered formations, the average count per cluster was 3.93 specimens, with a maximum of 32 individuals within one formation. 77.39% of sea anemones were actively hosting anemonefish. The cumulative *Heteractis magnifica* sea anemone coverage on the patch reef was 30.05m^2^ in May, 33.99m^2^ in July, and 34.49m^2^ by August. Other descriptive statistics for this reef region are presented in **Table 3**.

**Table 3.** Descriptive statistics for Heteractis magnifica monitored within the patch reef region of the research site at Village Reef, including formation and individual size, individual counts, and hosting status.

| Monitoring period | N | Formation size<br>(m<sup>2</sup>) |  | Individuals per cluster<br>(count) |  | Individual size<br>(m<sup>2</sup>) |  | Actively hosting | Solitary formations |
| --- | --- | --- | --- | --- | --- | --- | --- | --- | --- |
|  |  | Mean | SD | Mean | SD | Mean | SD | (%) | (%) |
| May | 177 | .170 | .220 | 3.71 | 4.43 | .048 | .029 | 74.01 | 45.20 |
| July | 195 | .174 | .251 | 4.25 | 5.19 | .041 | .023 | 80.00 | 48.21 |
| Aug | 203 | .170 | .258 | 3.84 | 5.07 | .047 | .037 | 77.83 | 51.23 |
| Total | 575 | .171 | .243 | 3.93 | 4.90 | .045 | .030 | 77.39 | 48.35 |

To test for significant effects of time on *Heteractis magnifica* abundance levels at Village Reef, Kruskal-Wallis analyses were run for formation size differences, individual sea anemone size differences, and cluster make-up differences over time. Results indicate no significant effect of time on sea anemone growth, including formation size, individual size and the counts of individuals clustered within a formation (*p* = .095, *p* = .290, and *p* = .309 respectively).

Moreover, to test for differences in formation size, individual size, and cluster makeup between the two reef habitats at Village Reef, Mann-Whitney U tests were performed. Results indicate that throughout the entire period of monitoring, a significant difference exists in the number of clustered individuals present within a formation between fringe and patch reef regions, where clustered formations contained significantly more individual specimens in the patch reef (see **Table 4**). No significant results were found for formation or individual sea anemone size throughout the monitoring period. Results did reveal a significant difference in cluster counts for May between the patch and fringe reef, with a mean count of 1.80 specimens per formation at the fringe reef, and a mean count of 3.71 in the patch reef (see **Table 4**). Formation cover was significantly different between reef habitats in July (see **Table 4**), with an average formation cover of .151m^2^ in the fringe reef, and .174m^2^ in the patch reef. The results provide partial support for our hypothesis of significant differences in *Heteractis magnifica* cover, size, and cluster make-up over time, as formation cove, and cluster counts did differ significantly between reef habitats within specific monitoring periods, although individual size estimates did not.

**Table 4.** Mann-Whitney U results for differences in Heteractis magnifica formation cover, size, and cluster make-up between the fringe and patch reef regions at Village Reef.

| Monitoring period | N | Test statistic | Stand. test statistic | p-value | S.E. |
| --- | --- | --- | --- | --- | --- |
| *Entire monitoring period* | 640 |  |  |  |  |
| Formation cover (m<sup>2</sup>) |  | 19511.500 | 0.583 | .074 | 1412.950 |
| Individual size (m<sup>2</sup>) |  | 16637.500 | -1.451 | .280 | 1412.950 |
| Cluster counts |  | 20873.000 | 1.670 | .048* | 1309.020 |
| *May* | 197 |  |  |  |  |
| Formation cover (m<sup>2</sup>) |  | 2145.500 | 1.556 | .356 | 241.680 |
| Individual size (m<sup>2</sup>) |  | 1680.500 | -0.370 | .060 | 241.680 |
| Cluster counts |  | 2287.000 | 2.310 | .011* | 223.770 |
| *July* | 219 |  |  |  |  |
| Formation cover (m<sup>2</sup>) |  | 2179.000 | -0.550 | .048* | 292.910 |
| Individual size (m<sup>2</sup>) |  | 1852.000 | -1.666 | .292 | 292.910 |
| Cluster counts |  | 2464.500 | 0.456 | .325 | 273.170 |
| *August* | 224 |  |  |  |  |
| Formation cover (m<sup>2</sup>) |  | 2185.500 | 0.177 | .430 | 282.718 |
| Individual size (m<sup>2</sup>) |  | 2045.500 | -0.304 | .381 | 282.719 |
| Cluster counts |  | 2211.000 | 0.306 | .380 | 259.920 |

Mann-Whitney U tests were performed to examine effects of hosting status on sea anemone formation cover, individual size, and cluster counts within the patch reef. Throughout the entire monitoring period, those *Heteractis magnifica* formations actively hosting anemonefish were larger than their non-hosting counterparts in formation cover, individual size, and cluster counts: *U* = 37181.500, *p* < .001 for formation cover; *U* = 31776.500, *p* =.044 for individual size; and *U* = 36596.500, *p* < .001 for cluster counts. When testing these effects for the individual monitoring periods, results remained significant for May: *U* = 5061.000, *p* < .001 for formation cover; *U* = 4208.000, *p* < .001 for individual size; and *U* = 4627.500, *p* < .001 for cluster counts. In July, only cluster counts had significant differences in hosting status: *U* = 3558.000, *p* =.040.

For August, no significant differences related to hosting status were detected. **Table 5** displays descriptive statistics for actively hosting and non-hosting *Heteractis magnifica*. As displayed in this table, sea anemones were larger when hosting, with a mean formation size difference of .159m^2^. Individual sizes were larger for actively hosting by .015m^2^ compared to non-hosting sea anemones. Moreover, cluster counts were larger for actively hosting formations, with 4.52 specimens on average in actively hosting formations, and 1.39 in non-hosting formations. These results are in keeping with the hypothesis of the study, although not all the expected effects regarding time and size estimates were found in the current study.

**Table 5.**
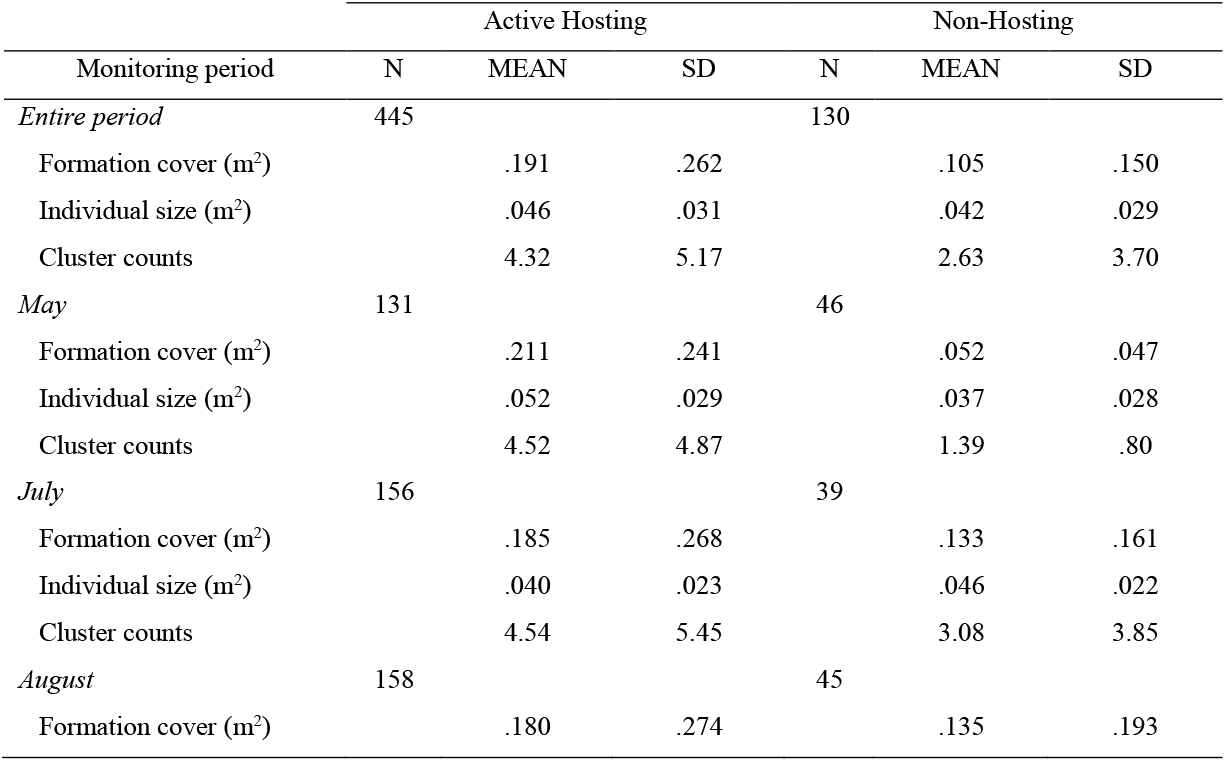
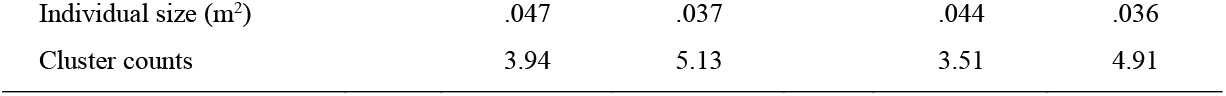
Descriptives for active- and non-hosting formations of Heteractis magnifica at the patch reef region at Village Reef.

Finally, to test for time effects on clustered versus solitary sea anemones, a ChiSquare test was run. Results revealed no significant effects of formation make-up over time, indicating that abundance levels of solitary or clustered formations did not differ significantly between the monitoring periods (*p* = .130), a result that contrasts our study hypothesis.

## DISCUSSION

The current study provided the first quantitative assessment of assemblages of *Heteractis magnifica* sea anemones located at Village Reef, Perhentian Kecil. As corals and sea anemones have been found to directly compete for suitable substrate and nutrients (Liu *et al*. 2009), understanding when and where sea anemones may outcompete corals when reef disturbances have occurred will help inform reef management and conservation programs around the Perhentian Islands. This study revealed several significant results, which provide preliminary insight into the population dynamics of the local distributions of *Heteractis magnifica*. The study yielded several significant effects of time on sea anemone formation cover, individual size, and cluster counts for the entire assemblage of *Heteractis magnifica* at Village Reef, with differential results depending on reef habitat and monitoring period. Within the deeper patch reef, larger cluster counts were found during May, and larger formation cover was seen during July. These findings are in support of literature on habitat requirements for sea anemones growth indicating seasonal and depth effects on sea anemone growth and asexual reproduction (Brolund *et al*. 2004; Elliot & Mariscal 2001; Holbrook & Schmitt 2005). Deeper regions and specific monitoring periods are associated with larger clusters containing more individuals at Village Reef.

A relationship of hosting status was present among the sea anemones at Village Reef. Actively hosting *Heteractis magnifica* were generally larger, contained more individuals within a cluster, and displayed increased formation cover compared to non-hosting sea anemones. This finding corroborates research proposing increased growth and asexual reproductive rates for actively hosting sea anemones and highlights the benefit of hosting (Brolund *et al*. 2004; Cleveland *et al*. 2011; Holbrook & Schmitt 2005; Porat & Chadwick-Furman 2004). However, significance of hosting and non-hosting formations changed throughout the monitoring period, with no significant differences found towards the pre monsoon period of August. This result highlights potential influences from seasonal changes including nutrient and temperature fluctuations and is supported by previous research indicating differential effects of cover, growth, and asexual reproduction related to seasonal effects (Holbrook & Schmitt 2005). Future research should focus on further detailing hostsymbiont dynamics to establish direct effects of hosting status on sea anemone growth.

In addition, with literature proposing a prominent effect of nutrient dynamics following the Northeast monsoon cycle, this topic deserves future attention. At the Perhentian Islands, nutrients levels reach peak concentrations in depth range 3-6 meters during the post monsoon phase in April (Mohamed *et al*. 2019). As such, expanding the monitoring period will enhance understanding of time related effects that remained outside the scope of the current study design. With both significant and insignificant effects, this study provides evidence of a dynamic change, including growth as the season approaches pre monsoon phase, and shrinkage in *Heteractis magnifica* size and cluster make-up between July and August. Future research should examine nutrient and temperature fluctuations at Village Reef, to establish how these are implicated in sea anemone growth. Such future directions are in line with studies on hosting sea anemones at other locations (Chomsky *et al*. 2004; Liu *et al*. 2009).

Despite our best efforts, there are some limitations to this study. Due to the limited monitoring period, assessment of time effects on the *Heteractis magnifica* population at Village Reef have limited applicability and would be strengthened by expanding the monitoring period. Ensuring the inclusion of more pre and post monsoon measurement will help to improve collective understanding of the impact of time relative to the monsoon on this specific assemblage of sea anemones. More so, indicators of nutrient level and temperature were not included in the current study due to a lack of reliable measuring instruments on location. As such, subsequent research would be enriched by additionally assessing these variables. More so, since solid evidence exists which highlights the influence of these variables on sea anemone distribution and growth (Adiana *et al*. 2014; Chomsky *et al*. 2004; Mohamed *et al*. 2019; Muller-Parker & Davy 2001), this aspect would be especially relevant as a future research direction.

Within this quantitative study of *Heteractis magnifica* abundance and assemblage dynamics at Village Reef, Perhentian Kecil, a first step has been made in better understanding *Heteractis magnifica* growth patterns locally. The current study adds insight to dynamics of aggregated sea anemones distributions and can play a valuable role in informing reef management and reef conservation programs. Restabilising the reefs on the Perhentian Islands is a critical task to ensure sustained viability of these reefs.

## CONCLUSION

In conclusion, the present study provided a first quantitative analyses of the local population of *Heteractis magnifica* sea anemones, including inspection of their aggregate forms, individual size, clustered counts, and hosting status at Village Reef, Perhentian Kecil, Malaysia throughout May, July, and August. Non -parametric testing revealed significant differences in cluster size per reef habitat region, and demonstrated the presence of larger sea anemone coverage, individual size estimates and cluster counts in actively hosting *Heteractis magnifica*. This study therefore contributes a first examination of the population of sea anemones present at Village Reef and helps to inform local reef management and conservation programs by providing valuable insight on *Heteractis magnifica* population dynamics.

